# Comorbid HIV and Cocaine Use Exacerbate Accelerated Brain Aging

**DOI:** 10.64898/2026.02.06.704435

**Authors:** Hong Gu, Betty Jo Salmeron, Danni Wang, Hong Lai, Nanyu Kuang, Hui Zheng, Shenghan Lai, Yihong Yang

**Author notes:** To whom correspondence may be addressed: Yihong Yang, Ph.D., Neuroimaging Research Branch, National Institute on Drug Abuse, National Institutes of Health, Baltimore, Maryland, USA.

## Abstract

**Background:** HIV and cocaine use (CU) each relate to cognitive deficits and brain abnormalities, yet their combined impact on brain aging remains unclear. This study examined how comorbid HIV and CU relate to brain aging and cognitive impairment.

**Methods:** We trained a morphometry-based brain-age model using harmonized Human Connectome Project–Aging data (HCP-A; n=725) with Gaussian Process Regression. The model was applied to an independent cohort with varying HIV/CU burden (HIV−/CU−, n=34; one disorder [HIV+/CU− or HIV−/CU+], n=72; HIV+/CU+, n=80). Brain age gap (BAG; predicted minus chronological age) was examined in relation to comorbidity burden and neurocognitive impairment (NCI; NIH Toolbox), adjusting for age, sex, education, depression, and image-quality indices. Analyses on SHapley Additive exPlanation (SHAP) values characterized network-wise feature-level contributions to brain age estimates.

**Results:** A dose-dependent effect of comorbidity burden on BAG was observed, with the HIV+/CU+ group showing the highest BAG. Greater BAG was associated with increased likelihood of NCI, and BAG partially mediated the relationship between comorbidity burden and NCI, with a stronger mediation effect in the two-disorder group than in the one-disorder group. Structural contributors to elevated BAG in the HIV/CU cohort included cortical thickness in the visual, ventral attention, and frontoparietal networks, and sulcal depth in the sensorimotor network.

**Conclusion:** Comorbid HIV/CU is linked to accelerated structural brain aging. BAG may reflect brain-level alterations underlying the association between comorbid HIV/CU and cognitive impairment, and may help identify network-specific targets for intervention.

## Introduction

Approximately 42-50% of people living with HIV (PLWH) are affected by HIV-associated neurocognitive disorders (HAND), a spectrum of conditions ranging from mild asymptomatic cognitive impairment to severe dementia (1). Although HIV-associated dementia has become rare with effective antiretroviral therapy (ART), milder forms of HAND remain prevalent, especially as the population of PLWH ages. These cognitive impairments, primarily involving deficits in executive functioning, psychomotor functioning, and information processing (2, 3), have been linked to structural and functional abnormalities in frontal, parietal, cingulate, and motor cortical regions (4–6).

Substance use, especially stimulant use, is also a risk for cognitive impairment (7, 8). Cocaine use (CU) is disproportionately prevalent in PLWH, with reported ever use of cocaine 4-5 times higher than those without HIV infection (9). Studies have shown that comorbid HIV and CU is associated with greater cognitive deficits (10–13) and additive structural brain abnormalities, particularly in parietal and occipital regions, where gray matter volume decrease is most pronounced in PLWH who use cocaine (14), although how HIV and CU interact is not yet entirely clear.

Structural brain changes that underpin these impairments are also hallmarks of aging. Brain ages are often estimated using machine learning models trained with structural neuroimaging data to quantify deviations from normative brain aging (15–17). The resulting brain age gap (BAG), defined as the difference between predicted and chronological age, has shown promise as a quantitative marker for brain health by identifying deviations from the normative aging trajectory (15, 18). More broadly, organ-specific biological age gaps, including the brain, can be estimated similarly and be utilized to predict chronic disease and mortality beyond chronological age, establishing their utility as integrative health indicators (19, 20).

In the context of HIV, prior studies have demonstrated elevated BAG in PLWH, even when HIV is effectively suppressed, and have linked increased BAG to worse cognitive performance (21). Similarly, increased brain aging has also been observed in individuals with cocaine use disorder, with larger BAG linked to longer duration of cocaine use (22). Yet, the combined impact of HIV and cocaine on brain age has not been systematically examined.

In this study, we built and cross-validated the brain age model using structural MRI data using a large healthy community sample from the Human Connectome Project-Aging (HCP-A) and then applied the model to predict brain age of participants in an HIV/CU cohort. To capture the cumulative burden of HIV and CU on brain aging, we grouped participants based on the number of neurobiological risk conditions: no disorder (HIV-/CU-), one disorder (HIV+/CU- or HIV-/CU+), and two disorders (HIV+/CU+). We aim to (1) characterize BAG profile across the three groups with graded neurotoxicity burden; (2) examine whether greater BAG relates to an increased risk of neurocognitive impairment (NCI); and (3) evaluate whether BAG mediates the relationship between cumulative disorder burden and NCI. We hypothesized that individuals with dual diagnoses (HIV+/CU+) would exhibit largest BAG, and that greater BAG would be associated with higher risk of cognitive impairment. Furthermore, we predicted that BAG would mediate the association between cumulative disorder burden and NCI, and that the specific networks and morphological features contributing to BAG may identify networks and features through which HIV and CU confer elevated risk for cognitive impairment, thereby identifying targets for prevention and treatment.

## Results

### Participant characteristics

The HIV/CU brain age prediction cohort was a subset of the parent Heart Study, a more than 20-year investigation supported by the National Institute on Drug Abuse examining the effects of HIV, ART exposure, and substance use on subclinical coronary artery disease (13, 23). The Heart Study recruited approximately 1500 adult men and women from the inner city of Baltimore, Maryland and was predominantly African American (93% of participants), reflecting the local demographic distribution. A total of 196 participants (aged 35-76.6 years, 84 females) from the HIV/CU comorbidity study completed neurocognitive assessments using NIH Toolbox Cognition Battery (NIHTB-CB) and underwent both structural and resting-state functional MRI (rsfMRI) scans. Although only structural data was used in brain age prediction, rsfMRI data was used to screen for excessive head motion, which can indicate compromised data quality in the structural scan acquired in the same session (24, 25). Participants were excluded if they failed FreeSurfer surface reconstruction (n=4) or showed excessive head motion during rsfMRI (n=6, defined as mean Euclidean displacement > 0.4 mm; see Supplementary Methods). This yielded a final sample of 186 participants for the brain age prediction analysis (Figure 1a).

**Figure 1.**
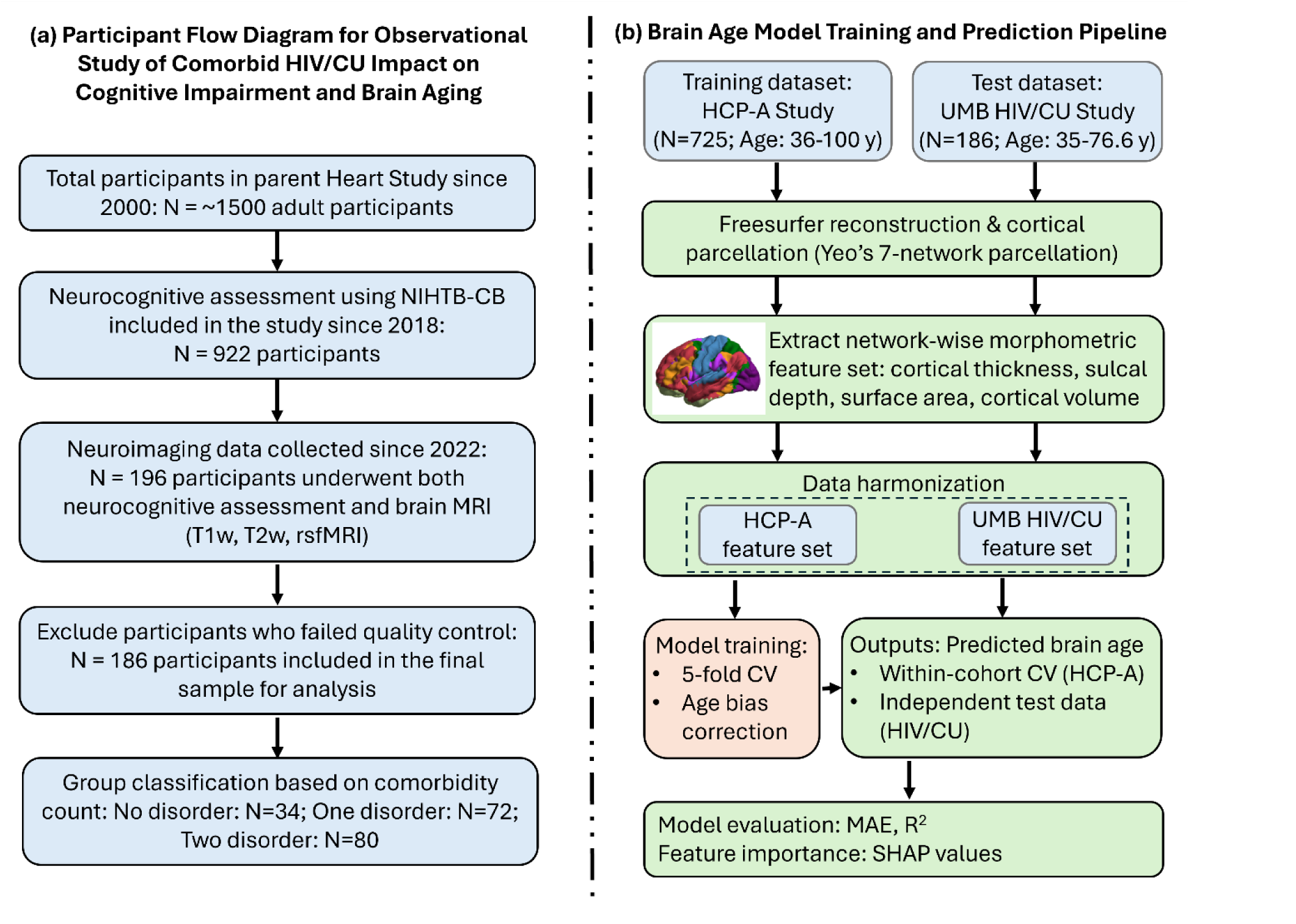
(a) Participant flow diagram for observational study of comorbid HIV/CU impact on cognitive impairment and brain aging; (b) Brain age model training and prediction pipeline. Network-wise mean cortical thickness, sulcal depth, total surface areas, and cortical volumes were calculated within visual (VIS), somatomotor (SMN), dorsal attention (DAN), ventral attention (VAN), limbic (LIM), frontoparietal (FPN), and default mode (DMN) network using Yeo’s 7-network parcellation atlas (29). Data harmonization was performed on the HCP-A and HIV/CU morphometric data to remove unwanted between-scanner variations from different acquisition sites and preserve biological variances associated with age, sex, and disease status (30). The harmonized HCP-A morphometric metrics were used to train the brain age prediction model using a nonlinear Gaussian Process Regression approach and then applied to the HIV/CU cohort to predict individual’s brain age. The prediction accuracy was evaluated with the mean absolute error (MAE) and the squared Pearson correlation (R^2^) between chronological age and predicted brain age. NIHTB-CB: NIH Toolbox Cognition Battery; HCP-A: Human Connectome Project Aging; CU: cocaine use; CV: cross validation; SHAP: Shapley Additive exPlanations.

To examine the combined impact of HIV infection and comorbid chronic cocaine use on cognitive impairment risk and brain aging, participants were categorized into 3 groups: 0 (HIV-/CU-), 1 (HIV+/CU- or HIV-/CU+), and 2 (HIV+/CU+), based on the number of disease conditions (HIV+ and/or CU+) present in each participant. To inspect if HIV+ and CU+ have distinct impact on clinical and cognitive outcomes, the 1-disorder group was further divided into two subgroups HIV+/CU- and HIV-/CU+. The demographic, clinical, and cognitive characteristics in each group are summarized in Table 1 and Table 2.

**Table 1.** Demographic and clinical characteristics of HIV/CU study participants (n=186)

| Characteristics | HIV-/CU-<br>(n=34) | HIV-/CU+<br>(n=39) | HIV+/CU-<br>(n=33) | HIV+/CU+<br>(n=80) | p-value |
| --- | --- | --- | --- | --- | --- |
|  | Disease count:0<br>(n=34) | Disease count:1<br>(n=72) |  | Disease count:2<br>(n=80) |  |
| Age <sup>a</sup> (years), mean (SD) | 56.9 (9.4) | 59.0 (8.3) | 56.3 (8.9) | 60.1 (6.6) | 0.066† |
|  | 56.9 (9.4) | 57.8 (8.6) |  | 60.1 (6.6) | 0.080† |
| Sex, female no. (%) | 18 (52.9) | 16 (41.0) | 10 (30.3) | 35 (43.8) | 0.31 |
|  | 18 (52.9) | 26 (36.1) |  | 35 (43.8) | 0.25 |
| African American, no.<br>(%) | 34 (100) | 34 (87.2) | 31 (93.9) | 80 (100) | 0.0024** |
|  | 34 (100) | 65 (90.3) |  | 80 (100) | 0.0017** |
| Education, high school<br>non-graduate no. (%) | 5 (14.7) | 7 (17.9) | 5 (15.2) | 20 (25) | 0.49 |
|  | 5 (14.7) | 12 (26.7) |  | 20 (25) | 0.31 |
| Zung depression scale,<br>mean (SD) | 40.9 (6.3) | 39.4 (9.0) | 41.2 (9.3) | 41.9 (9.2) | 0.56 |
|  | 40.9 (6.3) | 40.2 (9.1) |  | 41.9 (9.2) | 0.52 |

<sup>a</sup> Age: age at the MRI scan (years). \*\*: significance tested by Fisher's exact test.
Abbreviations: CU: Cocaine Use.

**Table 2.** Cognitive assessment with NIH toolbox cognition battery.

| NIH Toolbox Cognitive Measures | HIV-/CU-<br>(n=34) | HIV-/CU+<br>(n=39) | HIV+/CU-<br>(n=33) | HIV+/CU+<br>(n=80) | p-value <sup>a</sup> |
| --- | --- | --- | --- | --- | --- |
|  | Disease count:0<br>(n=34) | Disease count:1<br>(n=72) |  | Disease count:2<br>(n=80) |  |
| *Cognitive impaired <sup>b</sup>, no. (%)* | 6 (17.6) | 15 (38.5) | 13 (39.4) | 34 (42.5) | 0.039* |
|  | 6 (17.6) | 28 (38.9) |  | 34 (42.5) | 0.018* |
| *Total cognition composite* | 47.9 (11.9) | 44.8 (9.58) | 44.0 (12.0) | 42.9 (9.66) | 0.20 |
|  | 47.9 (11.9) | 44.5 (10.7) |  | 42.9 (9.66) | 0.14 |
| *Crystallized composite* | 52.2 (11.6) | 51.9 (10.0) | 49.5 (11.6) | 47.2 (9.44) | 0.044* |
|  | 52.2 (11.6) | 50.8 (10.8) |  | 47.2 (9.44) | 0.047* |
| Picture vocabulary | 46.9 (11.2) | 45.8 (10.8) | 45.1 (12.0) | 43.2 (9.75) | 0.36 |
|  | 46.9 (11.2) | 45.5 (11.3) |  | 43.2 (9.75) | 0.26 |
| Oral reading recognition | 57.0 (14.0) | 57.6 (10.7) | 53.8 (11.7) | 51.9 (11.1) | 0.031* |
|  | 57.0 (14.0) | 55.9 (11.3) |  | 51.9 (11.1) | 0.048* |
| *Fluid composite* | 44.2 (11.6) | 39.2 (10.7) | 40.2 (12.5) | 40.3 (10.4) | 0.28 |
|  | 44.2 (11.6) | 39.7 (11.5) |  | 40.3 (10.4) | 0.15 |
| Pattern comparison | 47.0 (13.7) | 39.5 (13.9) | 46.2 (14.2) | 44.4 (13.8) | 0.13 |
|  | 47.0 (13.7) | 42.6 (14.3) |  | 44.4 (13.8) | 0.39 |
| List sorting working memory | 45.1 (9.99) | 43.4 (9.96) | 39.8 (9.83) | 41.6 (9.98) | 0.10 |
|  | 45.1 (9.99) | 41.7 (9.99) |  | 41.6 (9.98) | 0.29 |
| Picture sequence memory | 48.3 (8.40) | 45.2 (8.01) | 47.6 (8.82) | 44.1 (9.02) | 0.18 |
|  | 48.3 (8.40) | 46.3 (8.42) |  | 44.1 (9.02) | 0.093† |
| Dimensional change card sort | 47.1 (14.6) | 45.0 (13.6) | 43.5 (13.9) | 44.7 (13.5) | 0.71 |
|  | 47.1 (14.6) | 44.3 (13.6) |  | 44.7 (13.5) | 0.64 |
| Flanker inhibitory control and attention | 44.0 (8.77) | 42.2 (10.3) | 41.6 (10.2) | 42.0 (8.99) | 0.68 |
|  | 44.0 (8.77) | 42.0 (10.2) |  | 42.0 (8.99) | 0.54 |

<sup>a</sup>Significance of group differences of all cognitive measures are assessed with age, sex, educational attainment, and Zung depression score controlled.
<sup>b</sup> Cognitive impairment was considered present if the fully adjusted T-scores was 1.0 SD below the mean of the fully adjusted standard scores involving at least two cognitive domains.

### Brain age prediction

Datasets acquired on 725 healthy participants (aged 36-100 years, 406 females) from the HCP-A Study Release 2.0 were used as a training cohort to build a brain age prediction model using nonlinear Gaussian Process Regression (Figure 1b) (26). Five-fold cross-validation revealed a high degree of correspondence between chronological age and predicted brain age within the HCP-A dataset (Supplementary Figure S1; mean absolute error MAE = 5.61 ± 0.25 years; squared Pearson correlation R^2^ = 0.83 ± 0.025 after age bias correction. Applying the trained model to the independent HIV/CU dataset, predicted brain age also showed high agreement with chronological age (Figure 2a; MAE = 5.71 years; R^2^ = 0.61 after age bias correction). The group-wise MAEs are 4.51, 4.93, and 6.92 years, and corresponding R^2^ values are 0.77, 0.63, and 0.51 for participants with 0, 1, and 2 HIV/CU conditions, respectively. With increasing HIV/CU burden, brain-age estimates showed progressively increasing deviation from normative brain aging, as reflected by increasing prediction error and reduced variance explained by the normative model.

**Figure 2.**
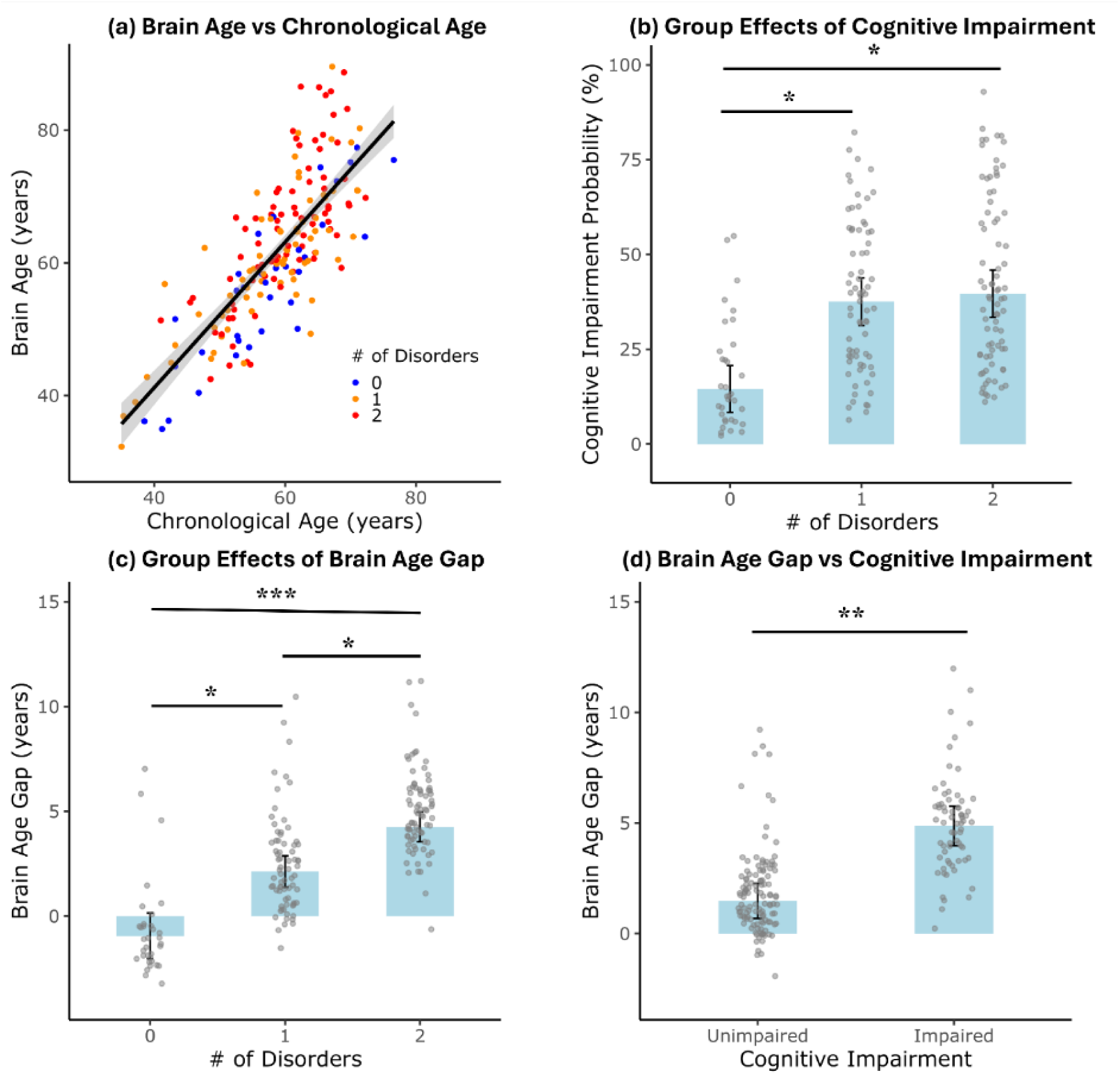
Group effects of HIV and cocaine use on brain age and cognitive impairment. (a) Relationship between predicted brain age and chronological age across participants, color-coded by number of disorders (0: healthy controls, 1: HIV+ or CU+ only, 2: comorbid HIV+/CU+); (b) Group effects on cognitive impairment probability showing higher impairment rates with disorders; (c) Group differences in brain age gap (BAG) across disorder groups, indicating greater BAG with increasing number of disorders; (d) Comparison of BAG between participants with and without cognitive impairment. Error bars represent the standard error of the mean. Significance level *: *p*<0.05; **: *p*<0.01; ***: *p*<0.001 (FDR corrected).

Brain age gap (BAG) was calculated as the difference between predicted brain age and chronological age. A positive BAG (predicted > chronological) may reflect accelerated, non-normative brain aging. Age, sex, educational attainment (high school non-graduate vs graduate), and the Zung self-rating depression scale (Zung score) were included in all statistical models testing for group differences and clinical associations with BAG. To control for motion-related bias in morphometric-derived BAG estimation, two proxy measures of image quality, the Euler number, a summary index of cortical surface reconstruction quality (27), and the Euclidean head motion index from rsfMRI, were also included in the regression models involving BAG, as head motion and image quality systematically influence cortical morphometric measures (24, 25) and head motion may covary with individual differences in cognitive functioning (28).

### Impact of comorbid disorders on cognitive impairment

Linear regression analyses were performed to assess potential differences in cognitive function and BAG between the HIV+/CU- and HIV-/CU+ subgroups. The results showed no significant differences in risk of cognitive impairment (*p* = .68) or BAG (*p* = .62) between the two subgroups (Supplementary Figure S2). These findings support combining the HIV+/CU- and HIV-/CU+ participants into a single subgroup in subsequent analysis.

Logistic regression revealed that greater HIV/CU disorder burden was associated with increased likelihood of NCI (χ²(2) = 7.13, *p* = .028). Model-adjusted predicted probabilities of NCI were 14.5% (95% CI: [6.0%, 31.1%]), 37.5% (95% CI: [26.2%, 50.4%]), and 39.7% (95% CI: [28.3%, 52.3%]) for individuals with 0, 1, or 2 HIV/CU disorders, respectively, after controlling for age, sex, education, Zung depressive score, and image quality proxy measures. Compared to individuals with no disorders, those with one disorder had 3.53 times greater odds (OR = 3.53, 95% CI [1.23, 11.46], *p* = .025), and those with two disorders had 3.87 times greater odds (OR = 3.87, 95% CI [1.36, 12.52], *p* = .016), while the difference in odds of NCI between one and two disorders was not significant (*p* = .81) (Figure 2b).

### Dose-dependent effect of comorbid disorders on brain age gap

A linear regression model was used to examine the association between the number of comorbid disorders and the morphometric BAG, controlling for age, sex, educational attainment, depressive symptom severity (Zung score), Euler number, and Euclidean head motion. A Type III ANOVA revealed a significant main effect of number of comorbid disorders (*F*(2, 177) = 10.83, *p* < .001).

Post-hoc pairwise comparisons (FDR-adjusted) showed that participants with one disorder had significantly higher BAG compared to those with no disorder (*mean difference* = 3.49 years, FDR adjusted *q* = .012), and those with two disorders had even greater BAG relative to those with no disorder (*mean difference* = 6.30 years, *q* < .001) (Figure 2c). Additionally, BAG was significantly higher in participants with two disorders than in those with only one (*mean difference* = 2.81 years, *q* = .012). These findings suggest a robust, cumulative association between comorbidity burden and accelerated brain aging.

### Cognitive impairment linked to elevated brain age gap

To examine the relationship between BAG and NCI, we conducted both linear and logistic regression analyses adjusting for age, sex, educational attainment, Zung depressive score, and image quality proxy measures (Euler number and head motion during fMRI acquisition). In the linear model with BAG as the outcome, a significant group effect was observed (*F*(1,178) = 9.03, *p* = 0.003), with cognitively impaired individuals showing higher BAGs (*mean difference* = 3.39 years), suggesting that cognitive decline is associated with accelerated brain aging (Figure 2d).

To assess whether BAG could predict NCI, a logistic regression was performed with NCI as the binary outcome and BAG as a predictor. BAG was significantly associated with the likelihood of NCI (*χ²*(1) = 9.79, *p* = 0.0018), with an odds ratio of 1.08 (95% CI: 1.03–1.15), indicating that each additional year increase in BAG raised the odds of cognitive impairment by 8%. These consistent findings across both regression approaches suggest that BAG not only reflects group differences in cognitive impairment but also serves as a meaningful predictor of individual NCI risk.

### Comorbidity-Related Cognitive Impairment Is Partially Mediated by Brain Age Gap

To assess whether the effect of the number of comorbid disorders on NCI was mediated by BAG, a structural equation model was constructed (31). The model included two dummy variables comparing individuals with 1 disorder (HIV+/CU- or HIV-/CU+) and 2 disorders (HIV+/CU+) to those with no disorders (HIV-/CU-), while controlling for age, sex, education, depressive symptoms (Zung score), and image quality (measured by Euler number and head motion).

As shown in Figure 3, both the indirect effect (β = 0.101, p = .022) and total effect (β = 0.298, p = .013) were significant for the 2-disorder group, whereas for 1-disorder group, only the total effect was significant (β = 0.274, *p* = .019), while the indirect effect was marginal (β = 0.055, *p* = .091). The mediation effect was marginally stronger in the 2-disorder group than in the 1-disorder group (*difference in indirect effect* = 0.046, *p* = .052), with no significant difference in total effects between the two groups (*p* = .85). These findings suggest that BAG may play an increasingly prominent mediation role in the relationship between comorbidity burden and NCI as the number of disorders increases, supporting the potential contribution of brain aging in comorbidity-related cognitive decline.

**Figure 3.**
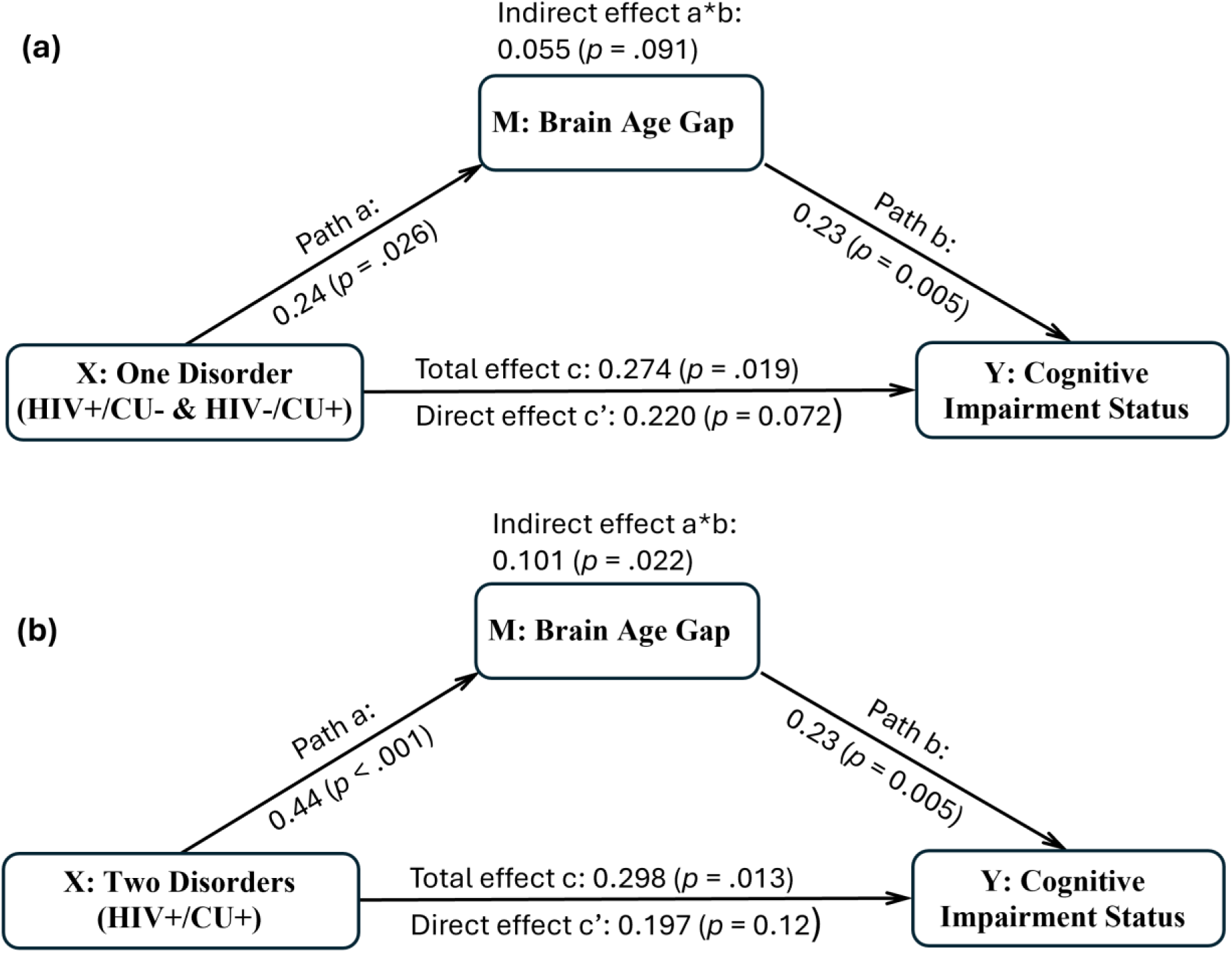
Structural equation model testing brain age gap (BAG) as a mediator of the association between HIV/cocaine use (CU) comorbidity and cognitive impairment status. Panel (a) depicts the contrast between individuals with one disorder (HIV+/CU- or HIV-/CU+) and those with no disorders (HIV-/CU-); and (b) shows the contrast between individuals with two disorders (HIV+/CU+) and those with no disorders. BAG significantly mediated the effect of comorbidity on cognitive impairment in the 2-disorder contrast and marginally mediated the effect in the 1-disorder contrast, with a marginally stronger indirect effect observed in the 2-disorder group than in the 1-disorder group (*p* = .052).

### Contributions of morphometric measures of brain networks to brain age prediction

#### Global patterns of feature importance across morphometric measures and networks

To quantify the contributions of network-wise morphometric measures to BAG prediction, SHapley Additive exPlanations (SHAP) values were estimated for each feature and participant. SHAP values provide local explanations of how each feature influences an individual’s predicted brain age, and global explanations by aggregating SHAP values across participants to assess overall feature importance (see Supplementary Methods) (32).

Global feature importance was quantified as the mean absolute SHAP value across participants, which index the magnitude of each feature’s contribution to BAG prediction irrespective of direction. Figure 4 summarizes the global feature importance of cortical thickness, sulcal depth, surface area, and cortical volume features across the Yeo’s seven canonical networks, which are visual (VIS), somatomotor (SMN), dorsal attention (DAN), ventral attention (VAN), limbic (LIM), frontoparietal (FPN), and default mode (DMN) networks, for both the HCP-A and HIV/CU cohorts, with networks ranked according to their importance in the HCP-A cohort.

**Figure 4.**
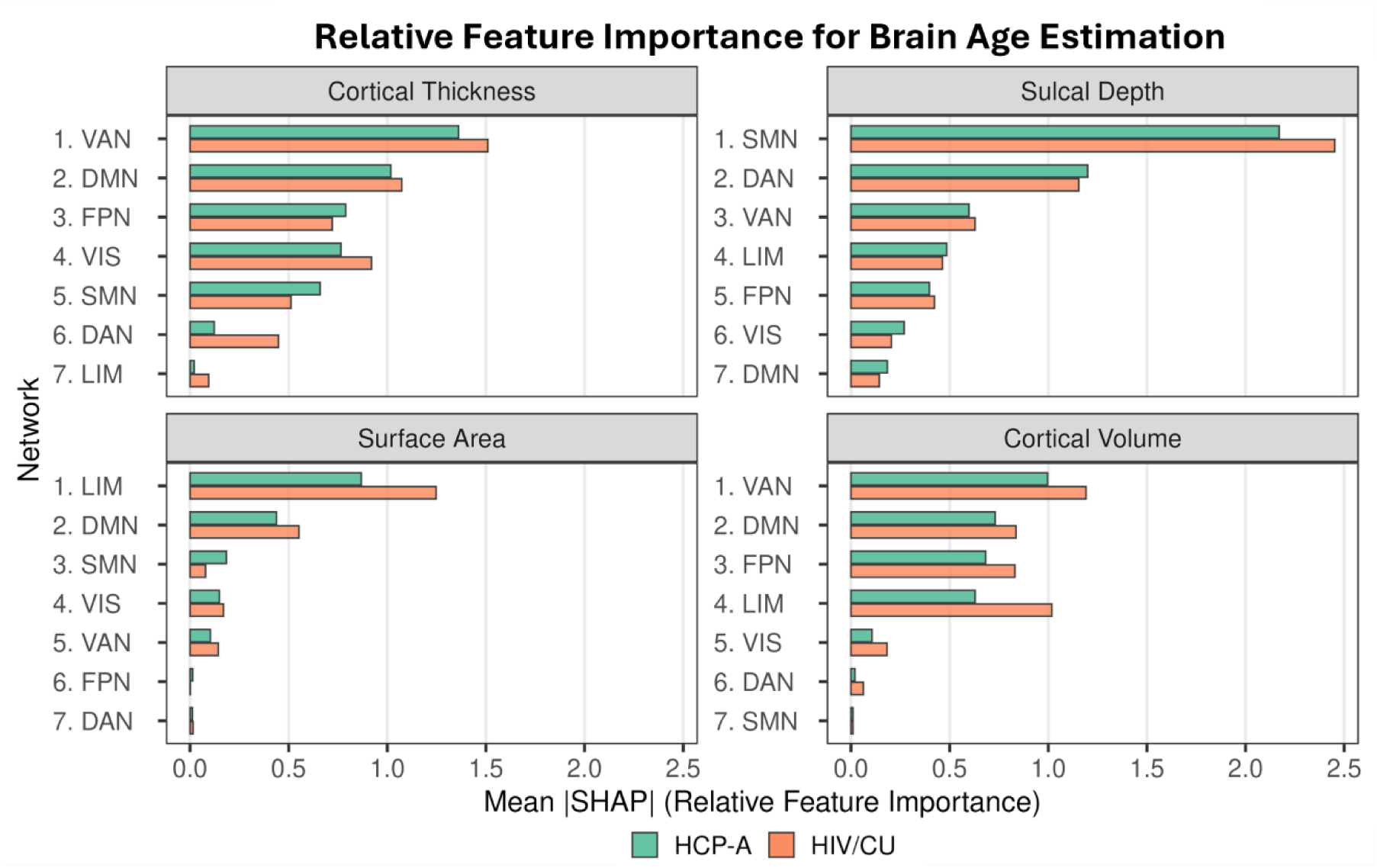
Relative overall feature importance of network-wise morphometric measures for brain age estimation. Horizontal bar plots show global SHAP feature importance, quantified as mean absolute SHAP values across participants, for four morphometric measures (cortical thickness, sulcal depth, surface area, and cortical volume) across seven large-scale brain networks. Within each morphometric measure, networks are ordered by descending importance in the HCP-A cohort, which serves as a normative reference. Side-by-side bars compare relative feature importance between the HCP-A (green) and HIV/CU (orange) cohorts. Higher values indicate a greater contribution of the corresponding network-wise morphometric feature to brain age estimation. Network abbreviations: VIS: visual network; SMN: somatomotor network; DAN: dorsal attention network; VAN: ventral attention network; LIM: limbic network; FPN: frontoparietal network; DMN: default mode network.

Both cohorts exhibited morphometric measure- and network-specific patterns of feature importance (Figure 4). For cortical thickness, feature importance was highest in the VAN and DMN, followed by FPN and VIS. In contrast, importance of sulcal depth was dominated by the SMN, with secondary contributions from the DAN. Surface area contributions were largely driven by the LIM, whereas cortical volume importance was primarily influenced by the VAN, DMN, and FPN. Across morphometric measures, the relative ordering of network importance was broadly similar between cohorts; however, the HIV/CU cohort exhibited systematically higher importance for limbic and association networks in several measures, suggesting cohort-specific shifts in the cortical features contributing to model predictions. These patterns indicate that while core aging-related signatures are shared between normative and comorbid populations, the relative contributions of specific network-wise morphometric measures are altered in the presence of HIV infection and substance use.

#### Comorbidity burden alters directional feature contributions in a network-specific and measure-specific manner

At the individual level, signed SHAP values quantify how much each morphometric feature increases or decreases predicted brain age relative to an age-matched reference baseline (constructed from k = 10 participants from the training set who were closest in age to each test participant) (33). Signed SHAP values thus provide a directional measure of feature influence on BAG prediction, with positive values indicating contributions toward older predicted age and negative values indicating contributions toward younger predicted age. To assess whether comorbidity burden (operationalized as disorder count: 0, 1, 2) modulates these directional contributions of network-wise morphometric features, linear mixed-effects (LME) models were fitted to signed SHAP values. Fixed effects included comorbidity burden (0-, 1-, and 2-disorders), network (VIS, SMN, DAN, VAN, LIM, FPN, and DMN), and morphometric measure type (cortical thickness, sulcal depth, surface area, and cortical volume), along with all higher-order interactions among these factors. Models controlled for age, sex, education, depression, and scan quality, and included random intercepts for participants to account for repeated measures and within-subject correlation.

The LME analysis revealed a significant comorbidity burden × network × measure type interaction (*F*(36, 4941) = 2.58, *p* < .001), indicating that the directional influence of morphometric features on brain age prediction depended jointly on comorbidity burden, network, and morphometric measure type. Significant lower-order interactions were also observed for comorbidity burden × network (*F*(12, 4941) = 2.52, *p* = .003), comorbidity burden × measure (*F*(6, 4941) = 5.07, *p* < .001), and network × measure (*F*(18, 4941) = 13.06, *p* < .001). Main effects of comorbidity burden (*F*(2, 177) = 10.86, *p* < .001), network (*F*(6, 4941) = 15.78, *p* < .001), and morphometric measure type (*F*(3, 4941) = 26.81, *p* < .001) were interpreted in the context of these interactions.

To unpack the significant three-way interaction, follow-up analyses tested comorbidity burden × network interactions separately for each morphometric measure.

##### Cortical thickness

The cortical thickness SHAP analysis revealed a significant comorbidity burden x network interaction (*F*(12, 4941) = 2.01, *p* = .020), with main effects of comorbidity burden (*F*(2,1354.52) = 11.60, *p* < .001) and network (*F*(6, 4941) = 8.11, *p* < .001). Post-hoc tests showed significant effects of comorbidity burden on thickness-related SHAP values in the VIS (*F*(2,5061.16) = 8.47, *p* = .00021, FDR corrected *q_FDR_* = .0015) and VAN (F(2,5061.16) = 6.84, *p* = .0011, *q_FDR_* = .0038), as well as a marginal effect in the FPN (F(2,5061.16) = 3.64, *p* = .026, *q_FDR_* = .061).

Pairwise comparisons indicated that higher comorbidity burden was associated with increased SHAP values in VIS (0 vs. 1: *p* = .046, *q_FDR_* = .046; 0 vs. 2: *p* = .0001, *q_FDR_* = .0002; 1 vs. 2: *p* = .014, *q_FDR_* = .021), VAN (0 vs. 1: *p* = .0016, *q_FDR_* = .0024; 0 vs. 2: *p* = .0004, *q_FDR_* = .0011; 1 vs. 2: *p* = .64), and FPN (0 vs. 1: *p* = .050, *q_FDR_* = .075; 0 vs. 2: *p* = .007, *q_FDR_* = .021; 1 vs. 2: *p* = .37), reflecting a greater positive contribution of cortical thickness to predicted brain age in these networks. Although effects in DMN did not survive correction, trends were consistent with higher SHAP values with increasing comorbidity burden (*F*(2,5061.16) = 2.80, *p* = .061, *q_FDR_*= .11; 0 vs. 1: *p* = .042, *q_FDR_* = .063; 0 vs. 2: *p* = .023, *q_FDR_* = .063; 1 vs. 2: *p* = .79). Collectively, these findings suggest that comorbidity burden modulates the directional contribution of cortical thickness to brain age prediction in a network-specific manner (Figure 5a).

**Figure 5:**
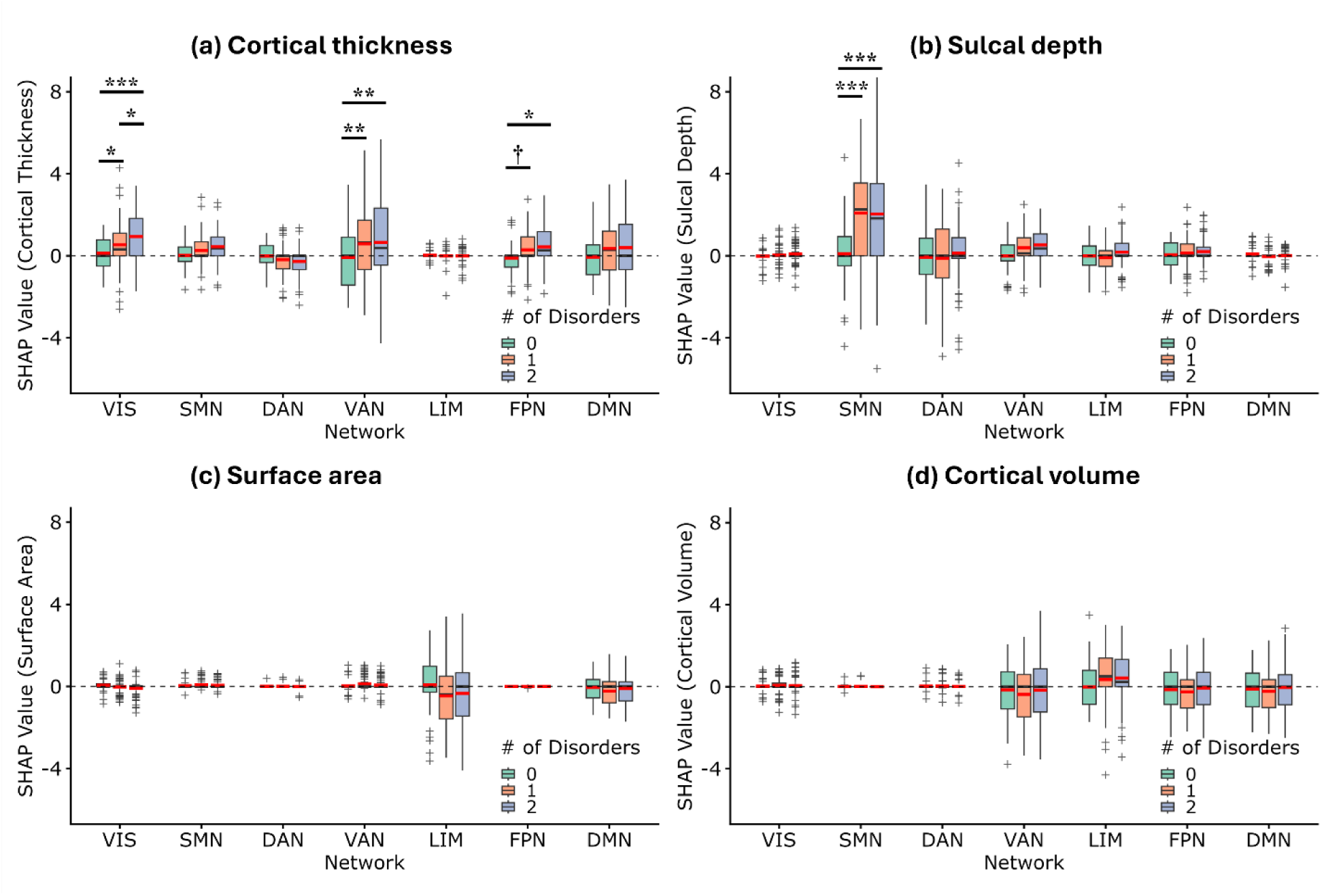
Comorbidity burden alters directional contributions of morphometric features to brain age prediction in a network- and measure-specific manner. Box-and-whisker plots show signed SHAP values for four morphometric measures (a) cortical thickness, (b) sulcal depth, (c) surface area, and (d) cortical volume across seven canonical brain networks. Boxes represent the interquartile range (25th–75th percentiles), inner black lines indicate the median, whiskers extend to the most extreme values within 1.5× the interquartile range, and “+” symbols denote outliers. Inner red lines indicate group means. The dashed horizontal line marks SHAP = 0, with positive values indicating features contributing to older predicted brain age and negative values indicating contributions toward younger predicted age. Statistical significance of pairwise differences across comorbidity groups within each network is FDR corrected: †: *p* < 0.10, *: *p* < 0.05, **: *p* < 0.01, ***: *p* < 0.001. Network abbreviations: VIS: visual network; SMN: somatomotor network; DAN: dorsal attention network; VAN: ventral attention network; LIM: limbic network; FPN: frontoparietal network; DMN: default-mode network.

##### Sulcal depth

For sulcal depth, a significant comorbidity burden x network interaction was observed (*F*(12, 4941) = 7.09, *p* < .0001), as well as main effects of comorbidity burden (*F*(2, 1354.52) = 14.84, *p* < .0001) and network (*F*(6, 4941) = 8.11, *p* < .0001). Follow-up analyses indicated that the interaction was driven primarily by the SMN, where individuals with one or two comorbid disorders showed significantly higher SHAP values than controls (*F*(2, 5061.16) = 51.32, *p* < .0001; both pairwise *p* < .0001). This pattern suggests increased directional positive influence of sulcal depth in the SMN on brain age prediction in participants with comorbidity burden (Figure 5b).

##### Surface area and cortical volumes

No significant comorbidity burden × network interactions were found for surface area or cortical volume (*all p* > .86), indicating that the directional contributions of these morphometric measures to brain age prediction did not vary with comorbidity across networks. A main effect of network was observed for cortical volume (*F*(6, 4941) = 4.00, *p* =.0005), reflecting network-dependent contributions averaged across comorbidity groups (Figure 5c-5d).

Together, these results indicate that comorbidity burden selectively alters the directional contributions of cortical thickness and sulcal depth to brain age prediction in a network-specific manner, whereas surface area and cortical volume reflect network effects that are not comorbidity dependent.

#### Morphometric validation of SHAP-identified comorbidity effects

To examine whether SHAP-identified differences in directional feature attribution corresponded to systematic variation in the underlying morphometric measures, follow-up linear models were fitted to cortical thickness and sulcal depth measures that showed significant comorbidity burden modulation in the SHAP analysis. Specifically, separate linear models were estimated to assess comorbidity effects of cortical thickness in the VIS, VAN, and FPN, and of sulcal depth in the SMN, controlling for age, sex, education, depression score, and scan quality.

Cortical thickness in the VIS network showed a significant main effect of comorbidity burden (*F*(2, 177) = 7.97, *p* = .00048, *q_FDR_* = .00097), and pairwise comparisons revealed progressively lower thickness with increasing comorbidity burden (0 vs. 1: *p* = .13; 0 vs. 2: *p* = .0008; 1 vs. 2: *p* = .009; Supplementary Figure S3a). A similar pattern of comorbidity effects was observed in the FPN (*F*(2, 177) = 4.01, *p* = .020, *q_FDR_* = .026), with pairwise tests indicating lower thickness in participants with higher comorbidity burden (0 vs. 1: *p* = .13; 0 vs. 2: *p* = .018; 1 vs. 2: *p* = .13; Supplementary Figure S3b). In contrast, cortical thickness in the VAN did not differ significantly by comorbidity burden (*F*(2, 177) = 1.83, *p* = .16; Supplementary Figure S3c), suggesting that SHAP attribution differences in this network may reflect multivariate dependencies not captured in univariate analyses.

For sulcal depth in the SMN, a robust main effect of comorbidity burden was identified (*F*(2, 177) = 9.10, *p* = .00017, *q_FDR_* = .00069). Post-hoc analyses indicated that individuals with one or two comorbid disorders exhibited significantly greater sulcal depth compared with those without comorbid disorders (0 vs. 1: *p* = .0003; 0 vs. 2: *p* = .0002; 1 vs. 2: *p* = .76; Supplementary Figure S3d), consistent with the strong comorbidity-related modulation observed in the SHAP analysis. These morphometric analyses partially validate the SHAP-based findings, demonstrating that comorbidity-related model attribution differences correspond to measurable structural differences in specific network and morphometric measure combinations. Notably, the discrepancy between SHAP and univariate morphometric results in regions such as the VAN highlights the advantage of SHAP framework: because SHAP values are derived from the full multivariate predictive model, they capture higher-order interactions and dependencies among features that may not be detectable in isolated univariate comparisons. Consequently, SHAP analysis provides enhanced sensitivity to complex, distributed patterns that contribute to brain age prediction.

## Discussion

In this study, we investigated the combined impact of comorbid HIV infection with cocaine use (HIV/CU) on cognitive impairment and brain aging, with a focus on BAG as a brain health biomarker. Specifically, we built and cross-validated a brain morphometry-based normative aging model trained on HCP-A data and applied the model to estimate brain ages of participants in an independent HIV/CU cohort, which was stratified by comorbidity burden into groups of 0-(HIV-/CU-), 1- (HIV+/CU- or HIV-/CU+), and 2-disorder (HIV+/CU+). We observed a dose-dependent increase in BAG with increasing comorbidity burden, in contrast to the plateauing of NCI, highlighting BAG’s sensitivity to cumulative neural burden. Cognitively impaired individuals had significantly higher BAGs, and greater BAG was associated with increased odds of NCI, reinforcing the notion that BAG captures subclinical neural decline beyond overt cognitive status.

### Brain age gap as a marker of accelerated aging and cognitive decline

Our findings demonstrate a dose-dependent increase in BAG with increasing comorbid HIV/CU burden. Specifically, BAG increases by approximately 3.5 years in individuals with one disorder and by more than 6 years when both conditions are present. This graded increase in BAG contrasts with the plateaued NCI risk beyond the presence of one disorder, suggesting that BAG captures ongoing neural stress even when clinical cognitive impairment appears to level off. Additionally, individuals with NCI exhibited significantly higher BAG, and each additional year of BAG increase was associated with increased odds of NCI, highlighting BAG’s sensitivity to cumulative neural burden and subclinical brain aging.

Converging evidence from mechanistic, structural, and functional domains corroborates this dose-dependent pattern of brain aging. Mechanistically, cocaine use may exacerbate HIV-induced neurotoxicity through neuroinflammatory and vascular pathways, including increased astrocyte and microglial activation, blood-brain barrier breakdown, and elevated viral replication, thereby placing synergistic stress on brain integrity (34–36). Structurally, PLWH who also use cocaine exhibit additive reductions in gray matter volume in the parietal and occipital cortices, exceeding the effects of either condition alone (14). Independent of HIV, cocaine use disorder has been associated with cortical thinning in association cortices and increased BAG, with greater BAG linked to longer duration of cocaine use, suggesting a substance-related acceleration of structural brain aging (22). Functionally, neuroimaging studies also aligns with this dose-response pattern: cocaine use attenuates compensatory prefrontal activation typically seen in PLWH during difficult choices in decision-making tasks, suggesting diminished neural reserve (37). In risk-processing tasks, only comorbid HIV+/CU+ participants exhibit hyperactivation in default-mode and cognitive control-related regions in response to increasing risk, indicative of compensatory but inefficient neural recruitment (38). Collectively, these findings align closely with our BAG results and underscore that comorbid HIV/CU condition imposes a greater impact on brain aging than either risk factor alone, even when overt cognitive impairment appears similar across comorbid groups.

Moreover, a recent large-scale study investigating the impact of comorbidities on brain aging in PLWH identified cardiovascular risk, hepatitis C co-infection, and socioeconomic adversity as primary contributors to elevated BAG (39). While acknowledging that variable selection techniques may overshadow related predictors, this study underscores the broader context in which non-HIV factors significantly influence brain aging. Within this context, our findings extend prior work by demonstrating that substance use, specifically cocaine, compounds HIV-related risk, reinforcing BAG’s potential as a dose-responsive biomarker of brain health that integrates multiple sources of neural stress.

### Mediation of comorbidity effects on cognition by brain aging

Mediation analysis using structural equation modeling indicated that BAG partially accounted for the relationship between comorbidity burden and neurocognitive impairment. The indirect effect was statistically significant in individuals with two comorbid conditions and marginally significant in those with one. Although total effects of comorbidity on NCI were significant in both contrasts, direct effects were only marginally significant after accounting for BAG, supporting the involvement of brain aging related processes in the pathway linking comorbidity burden to cognitive outcomes.

The mediation effect was marginally stronger in the 2-disorder group compared to the 1-disorder group, which may reflect greater engagement of brain aging related mechanisms as comorbidity burden increases. Nevertheless, a substantial portion of the comorbidity effect on NCI remains unexplained by BAG alone, suggesting the likely contribution of additional factors that did not contribute to BAG. Notably, the considerable overlap between network-wise morphometric measures with high importance on brain age prediction and those whose positive contribution increased with greater burden of HIV and CU suggests that these networks may be vulnerable to a wide range of neurobiological insults, further supporting the utility of BAG as an integrative marker of brain health. Given the cross-sectional design of the study, causal inferences cannot be drawn. Nonetheless, the findings support further investigation of brain aging as a potentially modifiable correlate of comorbidity related cognitive risk.

These results, in line with emerging evidence that BAG may mediate the effects of cognitive risk factors on cognitive outcomes (40), highlight brain aging as a potential therapeutic target. Interventions aimed at slowing brain aging through lifestyle modifications, vascular and metabolic health management, or neuroprotective treatments may help mitigate the neurocognitive consequences associated with high comorbidity burden and warrant further investigation in longitudinal studies.

### Network- and morphometric measure-specific alterations in brain age prediction

Using network-resolved SHAP value analyses, we identified morphometric measures of specific canonical brain networks whose contributions to the BAG were modulated by comorbidity burden. Notably, the overall pattern of network- and feature-level contributions to BAG was broadly similar between the HIV/CU cohort and the HCP-A cohort, in which BAG likely reflects the cumulative influence of non-diagnostic factors such as cardiovascular health, metabolic status, psychosocial stress, and physical fitness.

In some networks/features, there was substantial inter-individual variability in signed SHAP values, as reflected by wide interquartile ranges and extended whiskers in the box plots (Figure 5). In participants without comorbidities, median SHAP values often centered near zero, showing balanced positive and negative contributions to brain age prediction, which indicates that these morphometric features encode both vulnerability and resilience in healthy individuals. With increasing comorbidity burden, these distributions shifted upward to higher positive medians and reduced negative contributions, reflecting a predominance of aging-related contributions.

Within the HIV/CU cohort, cortical thickness in the VIS, VAN, and FPN, along with sulcal depth in the SMN, exhibited increasingly strong contribution to BAG with higher comorbidity burden. Functionally, the VAN supports stimulus-driven detection of salient events and rapid attentional reorienting, whereas the FPN underlies sustained attention, working memory, and flexible goal-directed control. These roles align with neuropsychological studies of HAND, which consistently identify attention, executive functioning, memory, and psychomotor speed as the most commonly affected cognitive domains in PLWH (2, 3, 41). Supporting this connection, prior neuroimaging work has shown associations between elevated BAG and poorer performance in information processing speed, executive function, and memory in HIV+ individuals (21).

Follow-up morphometric analyses confirmed that cortical thickness thinning in VIS and FPN was linked to increased comorbidity burden, consistent with extensive evidence that cortical thinning is a hallmark of structural brain aging and neurodegenerative processes. Reductions in cortical thickness in sensory and association networks has been linked to aging and disease progression in multiple populations (22, 42–44) and are frequently associated with declines in cognitive and sensorimotor functions, particularly in attention and executive domains that are commonly affected in HAND and substance use (14, 43, 45).

In contrast to prior literatures on brain aging and neurodegeneration, which typically report sulcal widening accompanied by reduced sulcal depth as a marker of global cortical atrophy (46, 47), we observed increased sulcal depth within the SMN as comorbidity burden increased. This apparent divergence likely reflects the region- and process-specific nature of sulcal morphology. Sulcal depth is by complex interactions among cortical thickness, underlying white matter integrity, and folding geometry, which may evolve heterogeneously across networks and disease stages. In primary sensorimotor cortex, which exhibits distinct developmental and aging trajectories relative to association cortex, localized deepening of sulci may arise from preferential thinning of gyral crowns and/or alterations in underlying white matter architecture. Accordingly, increased sulcal depth in the SMN does not necessarily contradict prior reports of global sulcal opening, but rather highlights the heterogeneity of cortical folding changes under cumulative disease burden.

Taken together, these morphometric findings indicate that comorbidity burden is associated not only with accelerated cortical thinning but also with alterations in cortical folding geometry within specific networks, reflecting complex, multi-factorial effects on cortical structure that are not fully captured by univariate measures alone. More broadly, our results align with growing evidence that cortical thickness and sulcal depth are sensitive and complementary markers of neurodegeneration and neuroinflammation. For instance, surface-based morphometry studies have revealed that both measures correlate with cognitive decline in Alzheimer’s disease and provide complementary insights (48), and that combining cortical thickness and sulcal depth improves discrimination of individuals with very mild Alzheimer’s disease, underscoring their potential diagnostic value (49).

### Implications and future directions

Individuals with HIV and a high comorbidity burden exhibited more rapid neurocognitive decline, even after accounting for HIV disease-related factors and demographic variables, underscoring comorbidity burden as a potent driver of cognitive deterioration over time (50). Our results provide additional evidence that comorbid conditions were associated with accelerated brain aging and increased neurocognitive risk, accompanied by distinct morphometric signatures that reflect network-level vulnerabilities. These vulnerabilities, identified through network-resolved SHAP values, appear to represent common targets of both neurotoxic insults and protective influences. The partial mediation on the relationship between HIV/CU comorbidity status and cognitive impairment by BAG further supports the potential of BAG as a biomarker for monitoring brain health and guiding intervention strategies in PLWH.

Longitudinal studies are needed to delineate the temporal trajectories of brain aging in the context of comorbidity and to determine whether interventions aiming at reducing comorbidity burden can slow down brain aging and preserve neurocognitive health. Furthermore, the differential impact of comorbidities on distinct morphometric measures and brain networks invites mechanistic investigation even as these vulnerabilities seem to be characteristic of the networks and measures rather than unique susceptibilities to HIV and CU. Integrating multimodal neuroimaging with molecular biomarkers holds promise for elucidating the biological mechanisms underlying comorbidity-driven brain aging (51). Understanding these complex interactions may facilitate the development of precision medicine strategies, where interventions are tailored to individual comorbidity profile and brain aging signatures.

### Individual and interactive effects of HIV and cocaine use

In this study, we used count of disorder as an index of comorbidity burden to examine the cumulative disease burden on NCI and brain aging. This analysis strategy simplified modeling and interpretation by using a single grouping variable. However, the effects of specific disorder type cannot be differentiated due to collapsing HIV-/CU+ and HIV+/CU- participants into 1-disorder group. To examine the individual and interactive effects of HIV and CU on cognitive impairment and brain aging, we performed additional analyses using the HIV × CU factorial models, controlling for the same confounding variables used in the main analysis.

A logistic regression model predicting NCI revealed significant main effects of both HIV status (χ² = 6.30, *p* = .012) and cocaine use (χ² = 4.14, *p* = .042). The interaction between HIV and CU showed a trend-level effect (χ² = 3.13, *p* = .077), exhibiting a semi-ordinal interaction pattern in which cocaine use increased NCI probability in HIV-individuals, but had limited additional impact in the HIV+ group (Supplementary Figure S4a). This pattern indicates that HIV status is the primary driver of cognitive impairment in this cohort, while cocaine use contributes primarily in the absence of HIV.

In the linear model predicting BAG, we observed significant main effects of both HIV status (χ² = 5.42, *p* = .020) and cocaine use (χ² = 4.29, *p* = .038), indicating that each condition was associated with an older predicted brain age relative to chronological age. Importantly, the HIV × CU interaction was not significant (*p* = .73), suggesting additive rather than synergistic effects. As shown in Supplementary Figure S4b, BAG increased with both HIV and cocaine use, with the highest BAG observed in HIV+/CU+ individuals.

### Limitations

While our study controlled for several confounding factors including age, sex, educational attainment, depressive symptoms, and imaging quality, potential residual confounding cannot be excluded. Confounding factors may include vascular and metabolic health, lifestyle behaviors, and inflammation, all of which have been linked to accelerated brain aging and cognitive decline in imaging studies (52–54). The cross-sectional design precludes causal inference and cannot discern whether accelerated brain aging is a cause or consequence of comorbidity-related cognitive decline.

The findings in this study should be interpreted in the context of the sample’s demographic profile. Given the predominantly African American composition of the cohort, the external validity to other racial/ethnic or geographic populations may be limited. However, the findings provide valuable insight into underrepresented groups often missing from neurocognitive research.

In summary, our findings demonstrate that the presence of comorbid conditions, particularly concurrent HIV infection and cocaine use, is associated with accelerated structural brain aging, as indicated by increased BAG. We identified morphometric vulnerabilities in the cortical thickness of the VIS, VAN, and FPN, along with the sulcal depth of the SMN, whose contributions to BAG were elevated in individuals with co-occurring HIV/CU condition. These network-specific structural alterations were similar to those in a generally healthy population but do align with the functional domains commonly impaired in HAND and the dose-dependent effect of comorbid HIV/CU on BAG is supported by mechanistic evidence that cocaine use may accelerate biological aging, as indicated by telomere shortening in PLWH who use cocaine (55). These converging findings suggest that BAG may serve as a biomarker of cumulative neurotoxic burden, especially in individuals with HIV and comorbid substance use.

## Methods

Detailed information on materials and methods is provided in Supplemental Methods.

### Sex as a biological variable

Both male and female participants were included in the HCP-A training dataset and the HIV/CU dataset. Biological sex was considered as a covariate in all statistical analyses examining brain age gap and neurocognitive outcomes.

### Statistics

Statistical analysis was conducted in R (version 4.5.0). Categorical parameters were summarized as count (proportions) and continuous variables were summarized as mean (SD). Linear and logistic regression analysis was performed to evaluate the relationships among comorbid HIV/CU disorder count, cognitive impairment, and morphometric BAG, controlling for age, sex, educational attainment (high school non-graduate vs. otherwise), the Zung self-rating depression scale (Zung score), and image quality indices (Euler number and Euclidean head motion index during rsfMRI). Specifically, participant head motion was quantified by average Euclidean distance between consecutive measurements across all functional scans. Using head motion index calculated during rsfMRI scan as a proxy for head motion during the structural scan is based on the observation that participant motion tended to be highly correlated across acquisitions within the same session and increased head motion during a functional sequence was associated with reduced morphometric estimates (24, 25).

Mediation analysis was performed to examine whether the impact of HIV/CU disease status on the risk of cognitive impairment was partially or fully explained by increased BAG using lavaan package (version 0.6.19) in R (31). The impact of disease status on feature importance, i.e. SHAP values, were investigated across brain networks and morphometric measures using linear mixed effects model (56).

### Study approval

The HIV/CU cohort data was collected under a protocol reviewed and approved by the institutional review board (IRB) at University of Maryland, Baltimore (13). All participants provided written informed consent prior to participation. The brain age prediction model training datasets are de-identified, publicly available data from the Human Connectome Project - Aging (HCP-A), which was collected under IRB approval with informed consent from all participants. Use of these data does not constitute human subjects research as defined under 45 CFR 46 and was exempt from additional IRB review.

### Data availability

The preprocessed HCP-Aging 2.0 Release data used in this report can be publicly accessed by qualified researchers from DOI: 10.15154/1520707 upon completion of a data use certification. The de-identified HIV/CU cohort data supporting the findings of this study may be made available upon reasonable request to the corresponding author, contingent upon institutional approval and data use agreements. Values for all data points in graphs are reported in the Supporting Data Values file.

## Author contributions

HG, BJS, SL and YY designed the research study; HG analyzed data and wrote the manuscript; DW, NK, and HZ helped design the project, provided analysis tool, and edited the manuscript; HL performed the experiment and acquired data; BJS, SL and YY contributed to the writing and editing of the manuscript, and supervised the study.

## Acknowledgements

This work utilized the computational resources of the NIH HPC Biowulf cluster (https://hpc.nih.gov).

## References

1. Wang Y, et al. Global prevalence and burden of HIV-associated neurocognitive disorder. Neurology. 2020;95(19):e2610–e2621.

2. Reger M, et al. A meta-analysis of the neuropsychological sequelae of HIV infection. Journal of the International Neuropsychological Society. 2002;8(3):410–424.

3. Deng L, et al. Association of HIV infection and cognitive impairment in older adults: A meta-analysis. Ageing Research Reviews. 2021;68:101310.

4. Gannon P, Khan MZ, Kolson DL. Current understanding of HIV-associated neurocognitive disorders pathogenesis. Current Opinion in Neurology. 2011;24(3):275.

5. Pfefferbaum A, et al. Accelerated and Premature Aging Characterizing Regional Cortical Volume Loss in Human Immunodeficiency Virus Infection: Contributions From Alcohol, Substance Use, and Hepatitis C Coinfection. Biological Psychiatry: Cognitive Neuroscience and Neuroimaging. 2018;3(10):844–859.

6. Wilson TW, et al. Multimodal neuroimaging evidence of alterations in cortical structure and function in HIV-infected older adults. Human Brain Mapping. 2015;36(3):897–910.

7. Potvin S, et al. Cocaine and Cognition: A Systematic Quantitative Review. Journal of Addiction Medicine. 2014;8(5):368.

8. Ramey T, Regier PS. Cognitive impairment in substance use disorders. CNS Spectrums. 2019;24(1):102–113.

9. Shiau S, et al. Patterns of drug use and HIV infection among adults in a nationally representative sample. Addictive Behaviors. 2017;68:39–44.

10. Avants SK, et al. Association between self-report of cognitive impairment, HIV status, and cocaine use in a sample of cocaine-dependent methadone-maintained patients. Addictive Behaviors. 1997;22(5):599–611.

11. Nigro SE, et al. Effects of cocaine and HIV on decision-making abilities. J Neurovirol. 2021;27(3):422–433.

12. Wakim K-M, et al. Assessing combinatorial effects of HIV infection and former cocaine dependence on cognitive control processes: A functional neuroimaging study of response inhibition. Neuropharmacology. 2022;203:108815.

13. Lai H, et al. Cocaine Use May Moderate the Associations of HIV and Female Sex with Neurocognitive Impairment in a Predominantly African American Population Disproportionately Impacted by HIV and Substance Use. AIDS Patient Care and STDs. 2023;37(5):243–252.

14. Bell RP, et al. Additive cortical gray matter deficits in people living with HIV who use cocaine. J Neurovirol. 2023;29(1):53–64.

15. Cole JH, et al. Brain age predicts mortality. Mol Psychiatry. 2018;23(5):1385–1392.

16. Cole JH, et al. Predicting brain age with deep learning from raw imaging data results in a reliable and heritable biomarker. NeuroImage. 2017;163:115–124.

17. Franke K, et al. Estimating the age of healthy subjects from T1-weighted MRI scans using kernel methods: Exploring the influence of various parameters. NeuroImage. 2010;50(3):883–892.

18. Kaufmann T, et al. Common brain disorders are associated with heritable patterns of apparent aging of the brain. Nat Neurosci. 2019;22(10):1617–1623.

19. Tian YE, et al. Heterogeneous aging across multiple organ systems and prediction of chronic disease and mortality. Nat Med. 2023;29(5):1221–1231.

20. Oh HS-H, et al. Plasma proteomics links brain and immune system aging with healthspan and longevity. Nat Med. 2025;31(8):2703–2711.

21. Cole JH, et al. Increased brain-predicted aging in treated HIV disease. Neurology. 2017;88(14):1349–1357.

22. Schinz D, et al. Lower cortical thickness and increased brain aging in adults with cocaine use disorder. Front Psychiatry. 2023;14. 10.3389/fpsyt.2023.1266770.

23. Lai H, et al. HIV and Low Omega-3 Levels May Heighten Hippocampal Volume Differences Between Men and Women With Substance Use. Brain, Behavior, & Immunity - Health. 2025;45:100988.

24. Pardoe HR, Kucharsky Hiess R, Kuzniecky R. Motion and morphometry in clinical and nonclinical populations. NeuroImage. 2016;135:177–185.

25. Savalia NK, et al. Motion-related artifacts in structural brain images revealed with independent estimates of in-scanner head motion. Human Brain Mapping. 2017;38(1):472–492.

26. Glasser MF, et al. The minimal preprocessing pipelines for the Human Connectome Project. NeuroImage. 2013;80:105–124.

27. Rosen AFG, et al. Quantitative assessment of structural image quality. NeuroImage. 2018;169:407–418.

28. Hausman HK, et al. The association between head motion during functional magnetic resonance imaging and executive functioning in older adults. Neuroimage: Reports. 2022;2(2):100085.

29. Thomas Yeo BT, et al. The organization of the human cerebral cortex estimated by intrinsic functional connectivity. Journal of Neurophysiology. 2011;106(3):1125–1165.

30. Fortin J-P, et al. Harmonization of cortical thickness measurements across scanners and sites. NeuroImage. 2018;167:104–120.

31. Rosseel Y. lavaan: An R Package for Structural Equation Modeling. Journal of Statistical Software. 2012;48:1–36.

32. Lundberg SM, et al. From local explanations to global understanding with explainable AI for trees. Nat Mach Intell. 2020;2(1):56–67.

33. Ball G, et al. Individual variation underlying brain age estimates in typical development. NeuroImage. 2021;235:118036.

34. Borrajo A, et al. Important role of microglia in HIV-1 associated neurocognitive disorders and the molecular pathways implicated in its pathogenesis. Annals of Medicine. 2021;53(1):43–69.

35. Buch S, et al. Cocaine and HIV-1 Interplay: Molecular Mechanisms of Action and Addiction. J Neuroimmune Pharmacol. 2011;6(4):503–515.

36. Dash S, et al. Impact of cocaine abuse on HIV pathogenesis. Front Microbiol. 2015;6. 10.3389/fmicb.2015.01111.

37. Meade CS, et al. Cocaine dependence modulates the effect of HIV infection on brain activation during intertemporal decision making. Drug and Alcohol Dependence. 2017;178:443–451.

38. Bell RP, et al. Neural sensitivity to risk in adults with co-occurring HIV infection and cocaine use disorder. Cogn Affect Behav Neurosci. 2020;20(4):859–872.

39. Petersen KJ, et al. Effects of clinical, comorbid, and social determinants of health on brain ageing in people with and without HIV: a retrospective case-control study. The Lancet HIV. 2023;10(4):e244–e253.

40. Tan WY, et al. Role of Brain Age Gap as a Mediator in the Relationship Between Cognitive Impairment Risk Factors and Cognition. Neurology. 2025;105(2):e213815.

41. Woods SP, et al. Cognitive Neuropsychology of HIV-Associated Neurocognitive Disorders. Neuropsychol Rev. 2009;19(2):152–168.

42. Salat DH, et al. Thinning of the Cerebral Cortex in Aging. Cereb Cortex. 2004;14(7):721–730.

43. Kallianpur KJ, et al. Regional Cortical Thinning Associated with Detectable Levels of HIV DNA. Cereb Cortex. 2012;22(9):2065–2075.

44. Makris N, et al. Cortical Thickness Abnormalities in Cocaine Addiction—A Reflection of Both Drug Use and a Pre-existing Disposition to Drug Abuse? Neuron. 2008;60(1):174–188.

45. Thompson PM, et al. Thinning of the cerebral cortex visualized in HIV/AIDS reflects CD4+ T lymphocyte decline. Proceedings of the National Academy of Sciences. 2005;102(43):15647–15652.

46. Kochunov P, et al. Age-related morphology trends of cortical sulci. Hum Brain Mapp. 2005;26(3):210–220.

47. Liu T, et al. The effects of age and sex on cortical sulci in the elderly. NeuroImage. 2010;51(1):19–27.

48. Sim Y, et al. Local gyrification index and sulcal depth as imaging markers of cognitive decline in Alzheimer’s disease. Front Aging Neurosci. 2025;17. 10.3389/fnagi.2025.1635861.

49. Ming J, et al. Integrated cortical structural marker for Alzheimer’s disease. Neurobiology of Aging. 2015;36:S53–S59.

50. Ellis RJ, et al. Higher Comorbidity Burden Predicts Worsening Neurocognitive Trajectories in People with Human Immunodeficiency Virus. Clin Infect Dis. 2022;74(8):1323–1328.

51. O’Connor EE, et al. Imaging of Brain Structural and Functional Effects in People With Human Immunodeficiency Virus. J Infect Dis. 2023;227(Supplement_1):S16–S29.

52. Angoff R, et al. Relations of Metabolic Health and Obesity to Brain Aging in Young to Middle-Aged Adults. Journal of the American Heart Association. 2022;11(6):e022107.

53. Bittner N, et al. When your brain looks older than expected: combined lifestyle risk and BrainAGE. Brain Struct Funct. 2021;226(3):621–645.

54. Franz CE, et al. Lifestyle and the aging brain: interactive effects of modifiable lifestyle behaviors and cognitive ability in men from midlife to old age. Neurobiology of Aging. 2021;108:80–89.

55. Lai S, et al. Cocaine use may induce telomere shortening in individuals with HIV infection. Progress in Neuro-Psychopharmacology and Biological Psychiatry. 2018;84:11–17.

56. Bates D, et al. Fitting Linear Mixed-Effects Models Using lme4. Journal of Statistical Software. 2015;67:1–48.

