## Supplemental Methods and Figures. for "Comorbid HIV and Cocaine Use Exacerbate Accelerated Brain Aging"

This file includes:

Supplemental Methods

Supplemental Figures

Supplemental References

### Supplemental Methods

#### Participants

##### *Lifespan Human Connectome Project Aging (HCP-A) Study*

Brain age estimation training data was provided by the Human Connectome Project Aging (HCP-A), funded by the National Institute on Aging under NIH grant U01AG052564 (Principal Investigators: David H. Salat, Susan Y. Bookheimer, and Marisa Terpstra). The HCP-A study was designed to discover brain aging patterns across healthy adulthood and aimed to enroll 1,500+ healthy adults representative of sex, race, ethnicity, and socio-economic status of the United States for an age range of 36-100+ years after excluding major diagnosed diseases (1). The preprocessed cross-sectional structural MRI datasets from 725 healthy participants (aged 36-100 years, 406 females) from the HCP-A Release 2.0 (DOI: 10.15154/1520707) were used as a training cohort to build a brain age prediction model.

##### *HIV and Cocaine Use on Cognitive Comorbidities Study*

Participants in this study are recruited from Baltimore city, Maryland under a protocol approved by the Institutional Review Boards at Johns Hopkins School of Medicine and University of Maryland, Baltimore (2). All participants provided written informed consent.

The study cohort included 1) HIV-infected and uninfected adults (≥18 years old) as determined by ELISA and confirmed by Western blot test; 2) chronic cocaine users, based on self-reported duration and frequency, and non-cocaine users as confirmed by a negative urine test for cocaine or benzoylecgonine. Exclusion criteria excluded: 1) significant neurological or psychiatric disorders; 2) contraindications to MRI; or 3) pregnancy, lactation, or childbearing potential.

#### Interview and medical chart review

Study participants underwent comprehensive interviews to collect information on sociodemographic characteristics, medical history, alcohol consumption, psychoactive substance use, cigarette smoking, and prescribed psychotropic medication use. The Zung Self-Rating Depression Scale was administered (3). For participants with HIV, additional data were gathered on HIV risk factors, duration of known HIV, and medication history including antiretroviral therapy (ART) use. Medical chart reviews were conducted to verify participants' medical history and prescribed medication information.

#### Neurocognitive Performance Assessment

Neurocognitive performance of participants in the HIV/Cocaine Use (CU) study was assessed with the app-administered NIH Toolbox Cognition Battery (NIHTB-CB), iOS version 1.20.2456 (www.nihtoolbox.org). The NIHTB-CB yields 7 individual test scores that measure 5 cognitive domains: language (Picture Vocabulary Test and Oral Reading Recognition Test), information processing speed (Pattern Comparison Processing Speed Test), working memory (List Sorting Working Memory Test), episodic memory (Picture Sequence Memory Test), and executive function (Dimensional Change Card Sort Test and Flanker Inhibitory Control and Attention Test). Three summary scores, Total Cognition Composite, Crystallized Composite, and Fluid Composite, were also provided based on the 7 individual test scores. Fully adjusted T-scores, which compares the score to those in the nationally representative normative sample after adjusting for age, sex, race, ethnicity, and educational attainment, were used to assess cognitive performance. Cognitive impairment was considered present if the fully adjusted T-scores was 1.0 standard deviation (SD) below the mean of the fully adjusted standard scores involving at least two cognitive domains, according to the revised research criteria for HIV-associated neurocognitive disorders (HAND) (4, 5).

#### MRI data acquisition

The HCP-A datasets were collected on Siemens Prisma 3T MRI scanners at four imaging sites. The T1w images were acquired with a multi-echo T1w MPRAGE sequence (resolution = 0.8×0.8×0.8 $mm^3$, TR/TI = 2500/1000 ms, TE = 1.8/3.6/5.4/7.2 ms, flip angle = 8°), and the final image was produced by the root-mean-square of the scans from all echoes. The T2w images were acquired with a T2w SPACE sequence (resolution = 0.8×0.8×0.8 $mm^3$, TR/TE = 3200/564 ms) (6).

For the HIV/CU datasets, T1w structural images were collected on a 3-T Siemens Prisma scanner (Erlangen, Germany) with a 3D MPRAGE sequence TE/TR/TI = 2.22/2500/1110 ms, flip angle = 8°, resolution = 0.8×0.8×0.8 $mm^3$. The T2w images were acquired with a T2w CAIPI sequence (resolution = 1×1×1 $mm^3$, TR/TE = 3200/349 ms) (7).

#### Surface Reconstruction and Quality Control

Surfaced-based brain morphometry was performed on anatomical T1w and T2w data using FreeSurfer 6.0 (8) for both HCP-A training datasets and HIV/CU test datasets to quantify vertex-wise cortical thickness, sulcal depth, surface area, and cortical volume measures. The released FreeSurfer data in HCP-A was quality controlled by visual inspection of all structural scans and FreeSurfer segmentation and surface outputs (9).

Quality control on the HIV/CU study dataset included visual check surface reconstruction quality and participants exclusion with large head motion. Motion during the rsfMRI scan was used as a proxy for head motion during the structural scan as an exclusion criterion. This criterion was based on the observation that participant motion tended to be highly correlated across acquisitions in the same session and increased head motion during a functional sequence was associated with reduced morphometric estimates (10, 11). Specifically, participant head motion was quantified by average Euclidean distance between consecutive measurements across all functional scans. Participants with mean Euclidean head motion larger than 0.4 mm were excluded from further analysis.

#### Brain Age Prediction

The HCP-A dataset was utilized to build and cross-validate the brain age prediction model, which was then applied to the independent HIV/CU dataset to predict individual's brain age.

*Feature extraction and data harmonization*: Mean cortical thickness and sulcal depth were calculated by 95% trimmed mean of vertex measures within visual network (VIS), somatomotor network (SMN), dorsal attention network (DAN), ventral attention network (VAN), limbic network (LIM), frontoparietal network (FPN), and default mode network (DMN) using Yeo's 7-network parcellation scheme (12). Surface area and cortical volume for each network were calculated as log-transformed sum of corresponding vertex measures in each network. ComBat, a batch-effect correction tool, was utilized to harmonize HCP-A and HIV/CU morphometric data to remove unwanted between-scanner variations from different acquisition sites and preserve biological variances associated with age, sex, and disease status (13). The disease status was coded as 1 if participant was HIV+ or CU+, otherwise was coded as 0. The harmonized data were then normalized to zero mean and unit variance.

*Model training and evaluation*: Model training and evaluation was performed using scikit-learn (1.3.0) in Python (3.10.8) (14). The 28 normalized and harmonized morphometric metrics (7 cortical thickness, 7 sulcal depth, 7 surface area, and 7 cortical volume) were used to train the brain age prediction model using a nonlinear Gaussian Process Regression (GPR) approach and then applied to the HIV/CU study cohort to predict individual's brain age. The prediction accuracy was evaluated with the mean absolute error (MAE) and the squared Pearson correlation ($R^2$) between chronological age and predicted brain age on the 5-fold cross-validation HCP-A datasets, which is stratified by scan site, as well as the independent HIV/CU dataset.

*Age bias correction*: Brain age estimation often exhibited an age-dependent bias, with an overestimation in younger participants and an underestimation in older participants than the mean age of the training dataset (15, 16). Bias correction for brain age estimation was applied to reduce this systematic estimation bias using the method proposed in (17).

#### Model Explanation and Feature Importance

SHapley Additive Explanations (SHAP) were employed to quantify the contribution of individual's morphometric measures of each network to the participant's brain age prediction (18). SHAP is based on cooperative game theory, attributing a model prediction (game outcome) to each input feature (player) by computing the average marginal contribution of that feature across all possible feature subsets. This framework ensures desirable properties such as consistency and local accuracy, making it suitable for interpreting complex, non-linear machine learning models.

SHAP values provide both local and global explanations. At the local level, each individual's SHAP value for a given morphometric feature reflects how much that feature increased or decreased the predicted brain age relative to a reference baseline. Global feature importance was obtained by averaging the absolute SHAP values across participants, which captures how strongly each feature influences predictions on average, regardless of direction. SHAP values are defined relative to a background reference set. If the reference set is sampled from the whole training set, the sum of SHAP values across all features of an individual equals the difference between his/her predicted brain age and the average brain age of the training dataset. In this study, a baseline reference set was comprised of *k* participants (k=10) from the training dataset with ages matched/closest to the age of each test participant. Using this age-matched background dataset, the SHAP values across all features summed to the individual's BAG.

#### Statistical Analysis

Statistical analysis was conducted in R (version 4.5.0). Categorical parameters were summarized as count (proportions) and continuous variables were summarized as mean (SD). Head motion can compromise structural image quality and bias morphometric estimation. Both FreeSurfer's Euler number and Euclidean head motion measure during fMRI scan from the same session showed associated with FreeSurfer-derived morphometric measures, especially cortical thickness, and were used as a proxy for structural image quality (10, 11, 19). To control for motion-related bias on morphometric-derived BAG estimation, Euler number and Euclidean head motion index during rsfMRI were included as covariates as imaging quality indices in all regression models. In addition, age, sex, education attainment (high school non-graduate vs. otherwise), the Zung self-rating depression scale (Zung score) were also controlled as potential confounding factors in the following regression models.

##### *Associations among disease status, cognitive impairment, and morphometric BAG*

The impact of disease status was investigated using a three-group categorization based on the number of comorbid conditions (HIV+ and/or CU+) each participant had: 0, 1, or 2 disorders.

*Disease impact on cognition*: Logistic regression analysis was performed to examine the impact of disease status on presence of cognitive impairment. Disease impact on the cognitive performance was investigated with linear regression analysis on 3 composite scores and 7 individual scores measured by NIHTB-CB.

*Disease impact on BAG*: The impact of disease status on BAG was examined by linear regression model with age, sex, education attainment, the Zung depression score, and imaging quality metrics (Euler number, and Euclidean head motion) as covariates.

*Association of increased BAG with cognitive impairment*: Both linear and logistic regression analysis were conducted to investigate if presence of NCI had impact on BAG as well as if increased BAG was associated with increased probability of cognitive impairment.

*Mediation effect of BAG*: Mediation analysis was performed to examine whether the impact of HIV/CU disease status on the risk of cognitive impairment was partially or fully explained by increased BAG using lavaan package (version 0.6.19) in R (20). Disease status was modeled using two mutually exclusive dummy variables representing individuals with 1 disorder (HIV+/CU- or HIV-/CU+) and 2 disorders (HIV+/CU+), with the no-disorder group as the reference. The outcome cognitive impairment status was modeled as an ordered factor to allow probit-based estimation. The model estimated indirect effects ($a \times b$), direct effects ($c'$), total effects, and the contrast of indirect effects between 2- vs 1- disorder. Models were estimated using diagonally weighted least squares (DWLS) with robust standard errors.

##### *Association between disease status and feature importance across networks*

To examine the impact of disease status on feature importance and to investigate if such impact varied across brain networks and morphometric measures, LME analysis was conducted on the SHAP values, which included fixed effects for disease count group, brain network, morphometric measure, and their interactions, with random intercepts for participant to account for repeated measures across networks and morphometric features, controlling for age, sex, education attainment, the Zung depression score, and imaging quality metrics (21).

#### Supplemental Figures

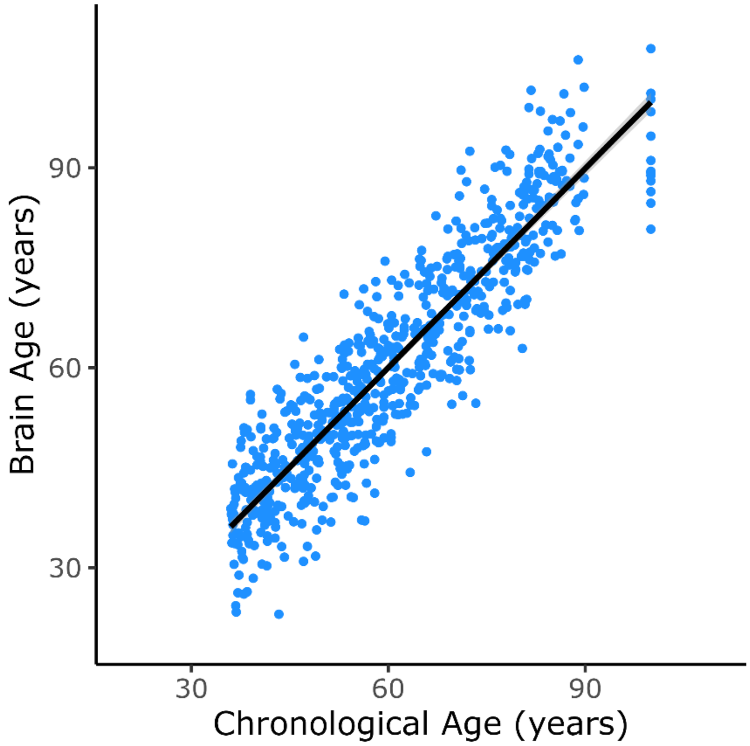


Supplementary Figure S1. Predicted brain age vs. participants' chronological age from Human Connectome Project Aging cohort with 5-fold cross validation.

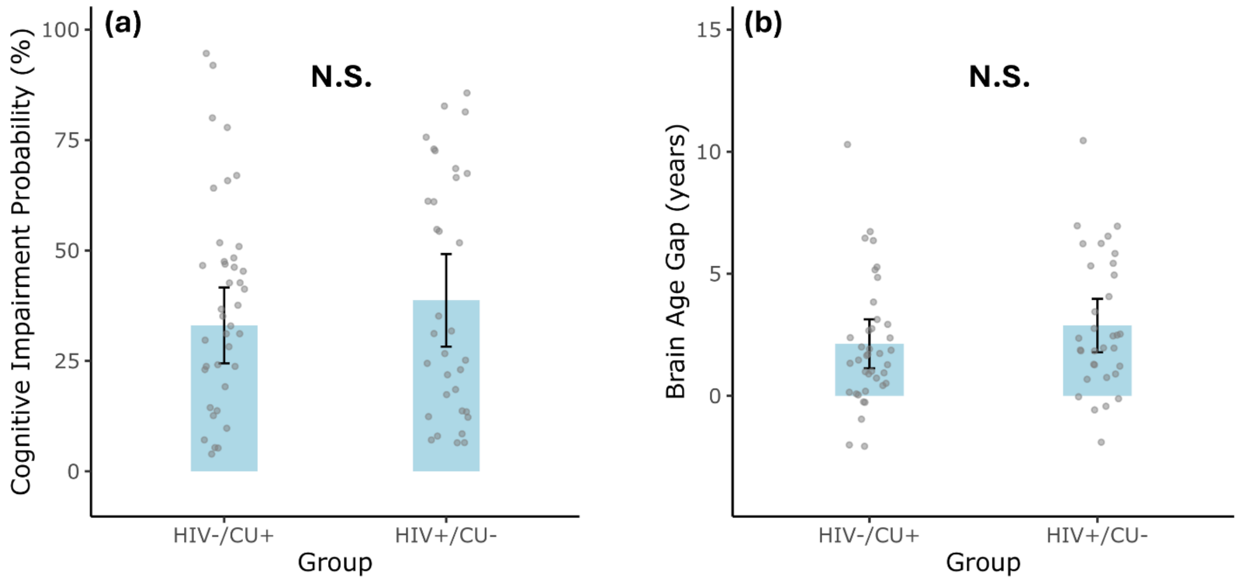


Supplementary Figure S2. No significant differences were found between the two HIV+/CU- and HIV-/CU+ subgroups in (a) risk of cognitive impairment and (b) brain age gap.

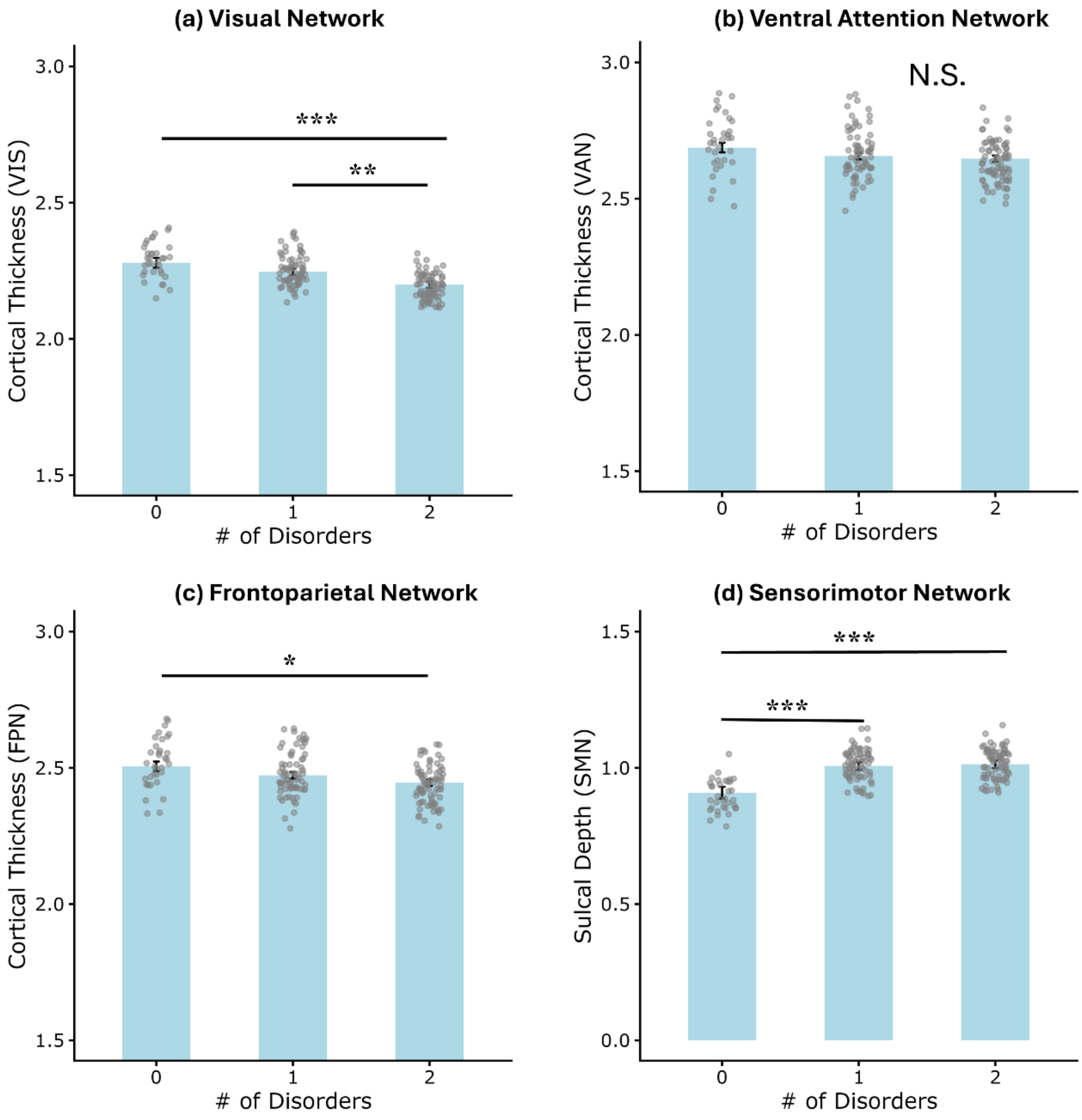


Supplementary Figure S3. Morphometric validation of SHAP-identified comorbidity effects on cortical measures. Bar plots with overlaid individual data points illustrate group differences in cortical thickness in the (a) visual network (VIS), (b) ventral attention network (VAN), (c) frontoparietal network (FPN), and (d) sulcal depth in the sensorimotor network (SMN), across levels of comorbidity burden (0: healthy controls, 1: HIV+ or CU+ only, 2: comorbid HIV+/CU+). Error bars represent the standard error of the mean. Significance level: *: $p<0.05$; **: $p<0.01$; ***: $p<0.001$; N.S.: non-significant differences (FDR corrected).

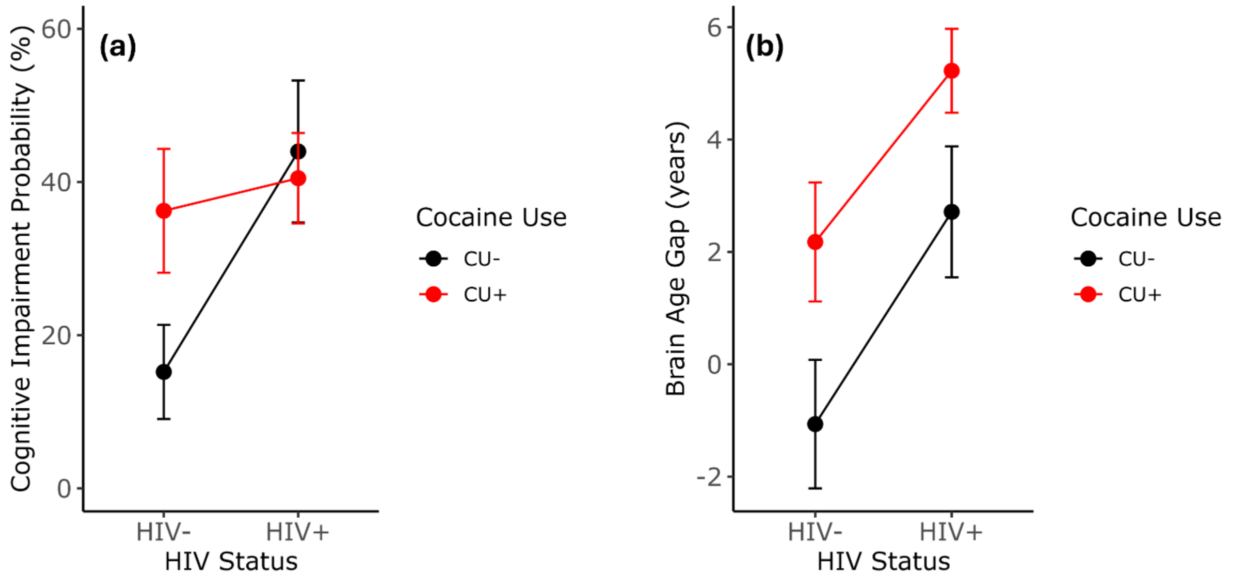


Supplementary Figure S4. Effects of HIV and cocaine use on cognitive impairment and brain age gap. (a) Estimated probability of cognitive impairment by HIV status and cocaine use. Both HIV and cocaine use were associated with higher impairment probability, with a trend-level interaction effect (p = .077). (b) Mean brain age gap (BAG) by HIV status and cocaine use. BAG was significantly elevated in both HIV+ and CU+ participants. Error bars represent standard error of the mean.

#### Supplemental References


1. Bookheimer SY, et al. The Lifespan Human Connectome Project in Aging: An overview. *NeuroImage*. 2019;185:335–348.

2. Lai H, et al. Cocaine Use May Moderate the Associations of HIV and Female Sex with Neurocognitive Impairment in a Predominantly African American Population Disproportionately Impacted by HIV and Substance Use. *AIDS Patient Care and STDs*. 2023;37(5):243–252.

3. Biggs JT, Wylie LT, Ziegler VE. Validity of the Zung Self-rating Depression Scale. *The British Journal of Psychiatry*. 1978;132(4):381–385.

4. Antinori A, et al. Updated research nosology for HIV-associated neurocognitive disorders. *Neurology*. 2007;69(18):1789–1799.

5. Clifford DB, Ances BM. HIV-associated neurocognitive disorder. *The Lancet Infectious Diseases*. 2013;13(11):976–986.

6. Harms MP, et al. Extending the Human Connectome Project across ages: Imaging protocols for the Lifespan Development and Aging projects. *NeuroImage*. 2018;183:972–984.

7. Lai H, et al. HIV and Low Omega-3 Levels May Heighten Hippocampal Volume Differences Between Men and Women With Substance Use. *Brain, Behavior, & Immunity - Health*. 2025;45:100988.

8. Fischl B, et al. Whole Brain Segmentation: Automated Labeling of Neuroanatomical Structures in the Human Brain. *Neuron*. 2002;33(3):341–355.

9. Marcus DS, et al. Human Connectome Project informatics: Quality control, database services, and data visualization. *NeuroImage*. 2013;80:202–219.

10. Pardoe HR, Kucharsky Hiess R, Kuzniecky R. Motion and morphometry in clinical and nonclinical populations. *NeuroImage*. 2016;135:177–185.

11. Savalia NK, et al. Motion-related artifacts in structural brain images revealed with independent estimates of in-scanner head motion. *Human Brain Mapping*. 2017;38(1):472–492.

12. Thomas Yeo BT, et al. The organization of the human cerebral cortex estimated by intrinsic functional connectivity. *Journal of Neurophysiology*. 2011;106(3):1125–1165.

13. Fortin J-P, et al. Harmonization of cortical thickness measurements across scanners and sites. *NeuroImage*. 2018;167:104–120.

14. Ball G, et al. Individual variation underlying brain age estimates in typical development. *NeuroImage*. 2021;235:118036.

15. Cole JH, et al. Brain age predicts mortality. *Mol Psychiatry*. 2018;23(5):1385–1392.

16. Le TT, et al. A Nonlinear Simulation Framework Supports Adjusting for Age When Analyzing BrainAGE. *Front Aging Neurosci*. 2018;10. https://doi.org/10.3389/fnagi.2018.00317.

17. Beheshti I, et al. Bias-adjustment in neuroimaging-based brain age frameworks: A robust scheme. *NeuroImage: Clinical*. 2019;24:102063.

18. Lundberg SM, et al. From local explanations to global understanding with explainable AI for trees. *Nat Mach Intell*. 2020;2(1):56–67.

19. Rosen AFG, et al. Quantitative assessment of structural image quality. *NeuroImage*. 2018;169:407–418.

20. Rosseel Y. lavaan: An R Package for Structural Equation Modeling. *Journal of Statistical Software*. 2012;48:1–36.

21. Bates D, et al. Fitting Linear Mixed-Effects Models Using lme4. *Journal of Statistical Software*. 2015;67:1–48.
